# Endothelial structure contributes to heterogeneity in brain capillary diameter

**DOI:** 10.1101/2023.04.26.538503

**Authors:** Sheridan M. Sargent, Stephanie K. Bonney, Yuandong Li, Stefan Stamenkovic, Marc Takeno, Vanessa Coelho-Santos, Andy Y. Shih

## Abstract

The high metabolic demand of brain tissue is supported by a constant supply of blood through dense microvascular networks. Capillaries are the smallest class of vessels and vary in diameter between ∼2 to 5 μm in the brain. This diameter range plays a significant role in the optimization of blood flow resistance, blood cell distribution, and oxygen extraction. The control of capillary diameter has largely been ascribed to pericyte contractility, but it remains unclear if endothelial wall architecture also contributes to capillary diameter heterogeneity. Here, we use public, large-scale volume electron microscopy data from mouse cortex (MICrONS Explorer, Cortical MM^3) to examine how endothelial cell number, endothelial cell thickness, and pericyte coverage relates to microvascular lumen size. We find that transitional vessels near the penetrating arteriole and ascending venule are composed of 2 to 5 interlocked endothelial cells, while the numerous capillary segments intervening these zones are composed of either 1 or 2 endothelial cells, with roughly equal proportions. The luminal area and diameter is on average slightly larger with capillary segments composed of 2 interlocked endothelial cells versus 1 endothelial cell. However, this difference is insufficient to explain the full range of capillary diameters seen in vivo. This suggests that both endothelial structure and other influences, such as pericyte tone, contribute to the basal diameter and optimized perfusion of brain capillaries.

## INTRODUCTION

The brain is a highly active and metabolically demanding organ. Capillary networks serve as the distribution network for oxygen and nutrient supply. Since the earliest in vivo imaging studies of the brain microcirculation, researchers have noticed a striking variance in blood flow and diameter among capillaries (1-4). Capillary segments of larger diameter tend to support higher flux of blood cells, and these vessels are spatially intermingled with thinner capillaries supporting lower flow (5, 6). While capillary flow does fluctuate on the timescale of seconds to minutes due to dynamic physiological processes such as vasomotion and neurovascular coupling (2, 7), these dynamics are built atop a relatively stable and heterogeneous pattern of capillary flow, set by capillary architecture and tone.

These findings raise the question of why heterogeneity in brain capillary flow is necessary. Recent studies suggest that heterogeneity creates reserve space for increased blood flow and tissue oxygenation during neural activity (*i*.*e*., functional hyperemia)(8). The idea builds on the capillary recruitment concept in peripheral tissues, such as muscle, where inactive tissue contains a proportion of non-flowing or very low-flow capillaries, which can be recruited to flow during activity (9). However, capillaries with no or very little flow is not ideal for brain function, given the high metabolic demand of neural tissue and lack of energy reserve. The system therefore establishes a continuous range of flow levels across all capillaries at baseline, with vessels lacking flow are very rare. This baseline state of heterogeneity transitions to more homogeneous flow among capillaries segments during increased brain activity. That is, low flux capillaries increase in flow, and high flux capillaries slightly decrease in flow (10, 11). Flow homogenization promotes more even distribution of oxygenated blood cells among capillaries, and slows their transit time to maximize oxygen extraction (8, 12).

Capillary diameter is a key determinant in setting basal flow heterogeneity. Capillaries normally range from ∼2-5 micrometers in diameter. Considering that red blood cells are on average 6 micrometers wide (white blood cells being larger), lumen diameter has a significant influence on flow resistance. Positive correlations between blood cell flux and capillary diameter have been measured in several studies, emphasizing the importance of capillary diameter regulation (6). Pericytes, the mural cells that line capillary networks, can regulate basal capillary diameter. Prior studies have shown that sustained optogenetic stimulation of pericytes can constrict capillaries in vivo, and conversely the ablation of pericytes leads to abnormal capillary dilation (6, 13, 14). Capillary pericytes possess some aspects of the contractile machinery expressed by arterial smooth muscle cells (15, 16). However, low to undetectable expression of key proteins such as alpha-smooth muscle actin confer very slow kinetics less likely to be involved in fast blood flow regulation, but adequate to support basal capillary flow heterogeneity (6). Single cell transcriptomic studies (17) and physiological studies both in vivo (18) and in vitro (19) also confirm that capillary pericytes express high levels of receptors for vasoconstrictive signaling, such as endothelin-1 type A receptors and thromboxane A2 receptors, potentially involved in basal capillary tone regulation.

While much has been learned about pericyte contributions to capillary tone, prior studies have not considered whether basic structural features of the capillary wall are sufficient to explain heterogeneity in brain capillary diameter. Here, we ask whether the number or thickness of endothelial cells constructing the capillary wall is related to the area and diameter of the capillary lumen. Addressing this question requires broad examination of capillary ultrastructure over the scale of entire capillary networks. With the availability of new large-scale volume EM resources such as the MICrONS Cortical MM^3 data set (20, 21), this question can now be rigorously tested.

## RESULTS

The MICrONS Cortical MM^3 data set contains numerous vertically-oriented penetrating arterioles and ascending venules. The dense microvascular networks intervening these routes of inflow and outflow (**Fig. 1A**). We categorized this vasculature into three zones (**Fig. 1B**): (*i*) arteriole-capillary transition (ACT; 4 branches from penetrating arteriole), (*ii*) capillary-venous transition (CVT; 4 branches from ascending venule), and (*ii*) capillaries (all intervening microvasculature between the ACT and CVT). The vascular lumen segmentation provided in the MICrONS explorer interface was used to navigate through the vascular architecture. We then specifically measured vascular attributes in the ACT, CVT, and capillary zones from 2D images of microvascular cross-sections (**Fig. 1C-E**).

**FIGURE 1.**
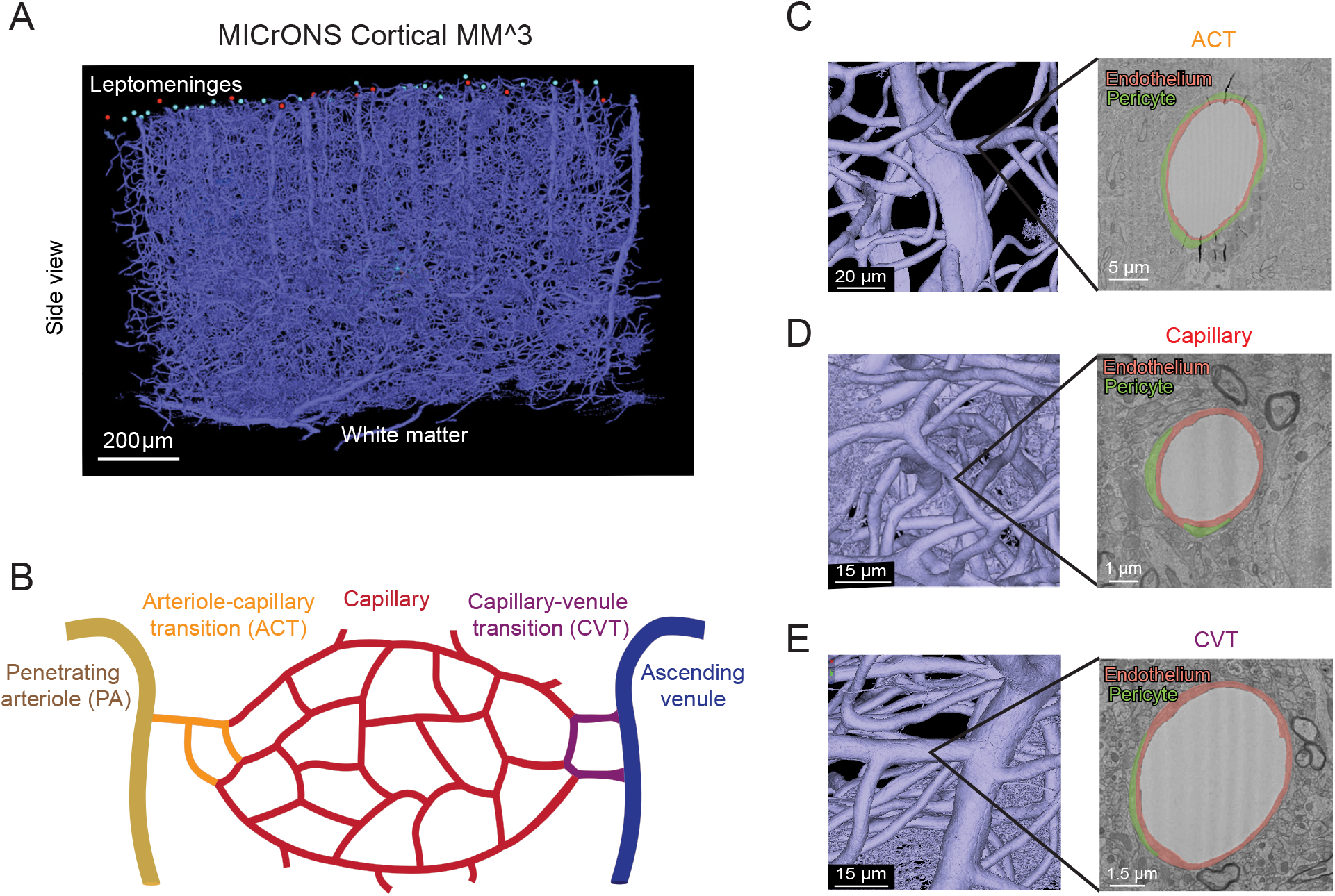
Different vascular zones can be examined within the MICrONS Cortical MM^3 data set. **(A)** Entire vascular segmentation within the Cortical MM^3 volume EM data set. **(B)** Schematic diagram showing different vascular zones as denoted in this study. **(C)** A penetrating arteriole with branching arteriole-capillary transition (ACT) vessel. A cross-section of the ACT vessel is shown, with endothelium and ensheathing pericyte highlighted in orange and green, respectively. The ACT ranges from 1-4 branch orders from the penetrating arteriole. **(D)** A capillary with the endothelium and pericyte processes highlighted. **(E)** A capillary-venule transition (CVT) zone vessel (left) and draining ascending venule with the endothelium and pericyte processes highlighted. For this study, we denoted the CVT range as 1-4 branch orders from the ascending venule.

We quantified lumen area, lumen diameter, endothelial area, and endothelial thickness of individual vessels across the three microvascular zones (**Fig. 2A-D**). Microvessels were sampled across all cortical layers (and superficial corpus callosum) and then pooled to gain an overall view of their characteristics. Lumen area and diameter of ACT and CVT zones were larger than capillaries (**Fig. 2A,B,E,F**). A broad range of diameters were measured in each microvascular zone. As expected, the area of the endothelium was greater with larger diameter vessels (**Fig. 2C,G**). Interestingly, endothelial thickness was significantly larger in ACT and CVT zones, compared to capillaries (**Fig. 2D,H**).

**FIGURE 2.**
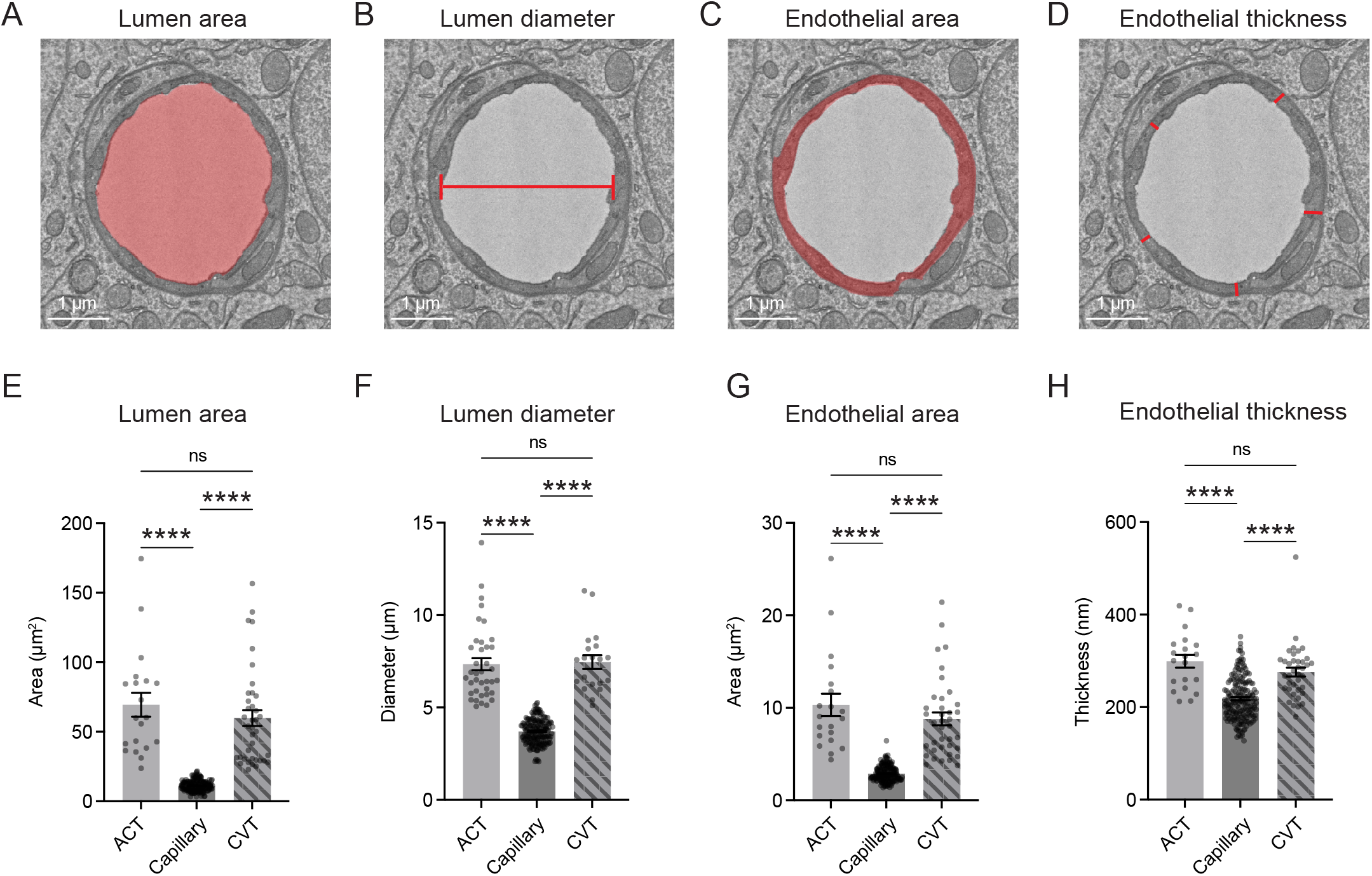
Vessel characteristics across capillary and transitional zones. **(A)** Measurement of vessel lumen area, as indicated by region of red shading on representative image of capillary. **(B)** Measurement of lumen diameter. Capillary lumen diameter was extrapolated from the lumen area. For ACT and CVT zones, lumen diameter was the length of the minor axis given their occasional oval shapes. **(C)** Measurement of endothelial area, as indicated by red shading. **(D)** Endothelium thickness, as recorded from 5 locations and averaged per vessel cross-section. **(E)** Comparison of lumen area across microvascular zones. Kruskal-Wallis test ****p<0.0001. Dunn’s multiple comparisons test showed no difference between ACT and CVT, p>0.99. ACT vs Capillary ****p<0.0001. Capillary vs CVT ****p<0.0001. **(F)** Comparison of lumen diameter across microvascular zones. Kruskal-Wallis test ****p<0.0001. Dunn’s multiple comparisons test showed no difference between ACT and CVT, p>0.99. ACT vs Capillary ****p<0.0001. Capillary vs CVT ****p<0.0001. **(G)** Comparison of endothelium area across microvascular zones. Kruskal-Wallis test ****p<0.0001. Dunn’s multiple comparisons test showed no difference between ACT and CVT, p>0.99. ACT vs Capillary ****p<0.0001. Capillary vs CVT ****p<0.0001. **(H)** Comparison of endothelial thickness across microvascular zones. Kruskal-Wallis test ****p<0.0001. Dunn’s multiple comparisons test showed no difference between ACT and CVT, p>0.99. ACT vs Capillary ****p<0.0001. Capillary vs CVT ****p<0.0001. All data shown as mean ± SEM.

To determine whether capillary diameter heterogeneity is influenced by the number of endothelial cells in the vessel wall, each capillary cross section was examined for tight junction (TJ) number by 3 independent raters (SMS, SKB, VCS). A capillary with a single TJ is composed of a single endothelial cell wrapping and connecting with itself (**Fig. 3A**). A capillary with 2 TJs is composed of two endothelial cells interlocking to form the vessel wall (**Fig. 3B**). The capillary zone contained predominantly capillaries with 1 or 2 TJs, with roughly equal proportions (**Fig. 3C**). Capillaries with 3 and 4 TJs, indicating 3 and 4 endothelial cells respectively, were also observed on rare occasions (3 out of 185 capillaries examined). In contrast, these multi-TJ vessels were common in the ACT and CVT zones, with up to 5 or 6 TJs per vessel observed (**Supplementary Fig. 1**).

**FIGURE 3.**
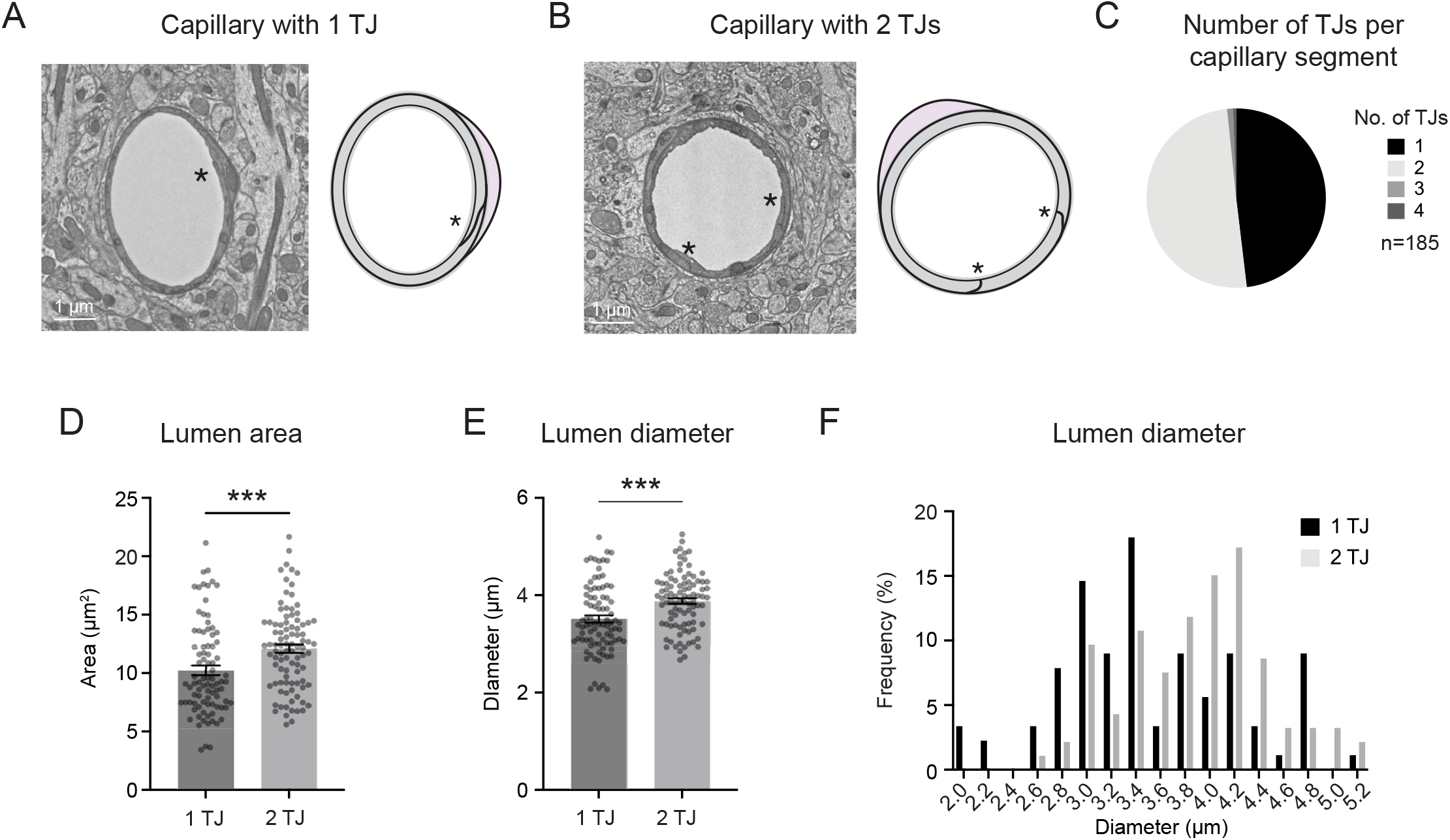
Influence of endothelial cell number on capillary size. **(A)** Representative image and schematic of capillary with one tight junction (1 TJ), and therefore 1 endothelial cell. **(B)** Representative image and schematic of capillary with two tight junctions (2 TJ), and therefore two interlocked endothelial cells in the vessel wall. **(C)** Distribution of TJ number across 185 capillaries. Our sampled group had 89 (48.11%) capillaries with 1 TJ, and 93 (50.27%) capillaries with 2 TJs. Only 2 capillaries (1.08%) were found with 3 TJs and 1 capillary (0.54%) found with 4 TJs. **(D)** Comparison of lumen area between capillaries with 1 and 2 TJs. Unpaired t test (two-sided), t(178)=3.357;***p=0.0010. N=89 capillaries with 1 TJ, n=93 capillaries with 2 TJ. Data shown as mean ± SEM. **(E)** Comparison of lumen diameter between capillaries with 1 and 2 TJs. Unpaired t test (two-sided), t(170.3)=3.854;***p=0.0002. N=89 capillaries with 1 TJ, n=93 capillaries with 2 TJ. Data shown as mean ± SEM. **(F)** Frequency distribution of lumen diameter of capillaries with 1 and 2 TJs.

We compared vascular attributes between capillaries with 1 or 2 TJs. Lumen area and diameter were on average significantly larger with 2 TJ capillaries. However, the contribution of endothelial number to capillary lumen size was modest (**Fig. 3D,E**). Average capillary areas and diameter were only 2 μm^2^ and 0.4 μm larger with 2 TJs compared to a single TJ, respectively (**Fig. 3D,E**). In contrast, the range in capillary area (∼3-21 μm^2^) and diameter (∼2-5 μm) was broad and overlapped heavily between the two groups (**Fig. 3F**). This suggests that endothelial number contributes to capillary diameter, but other factors beyond cell number, such as pericyte tone, must also contribute to capillary diameter heterogeneity.

To further verify whether endothelial cell number influences capillary lumen area, we examined the upper and lower extremes of lumen area within the capillary group. We separated the smallest 25% of lumen areas (lower) and the largest 25% of lumen areas (upper) and compared the distribution of endothelial TJs numbers in these groups (**Supplementary Fig. 2**). Vessels in the lower 25% group were composed mostly of those with 1 TJ, while the upper 25% were predominantly vessels with 2 TJs, and also contained the rare vessels with 3 and 4 TJs.

We next examined how other attributes of the endothelium related to lumen size. As expected, the area of the endothelial cross-section was greater with the larger lumen areas of 2 TJ capillaries (**Fig. 4A**). We also considered if distention of endothelium was necessary to create larger diameter capillaries, *i*.*e*., whether larger capillaries have thinner walls due to cell stretching. Instead, we found that larger capillaries exhibited thicker endothelial walls, and that there was an overall positive relationship between lumen area/diameter and endothelial thickness (**Fig. 4B-D**).

**FIGURE 4.**
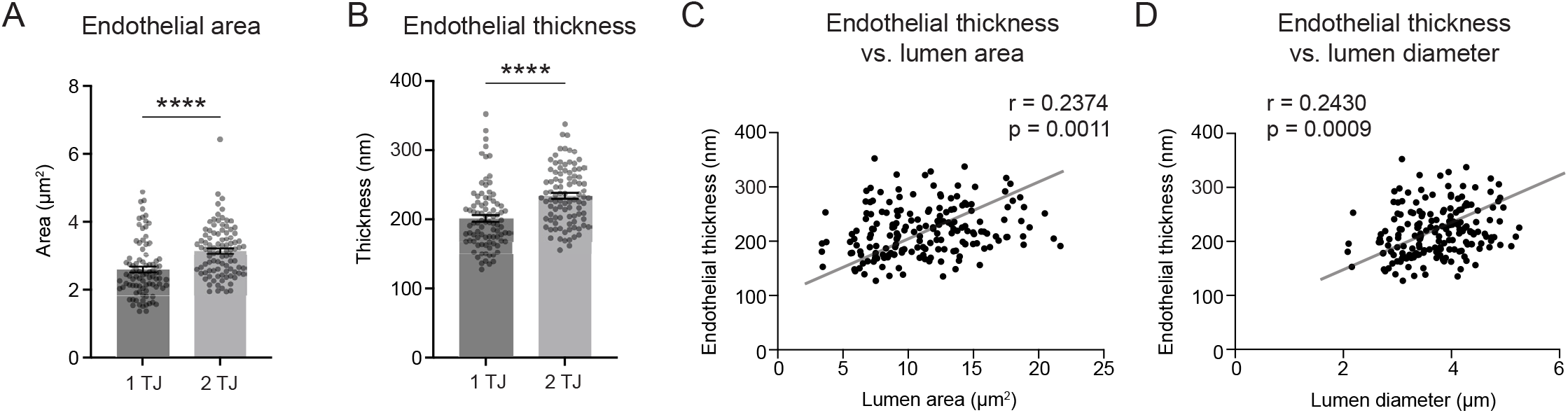
Relationship between endothelial area and thickness on capillary size. **(A)**Comparison of endothelial area between 1 TJ and 2 TJ capillaries. Unpaired t test (two-sided), t(180)=4.773;****p<0.0001. N=89 capillaries with 1 TJ, n=93 capillaries with 2 TJ. **(B)** Comparison of endothelial thickness between 1 TJ and 2 TJ capillaries. Unpaired t test (two-sided), t(180)=5.076;****p<0.0001. N=89 capillaries with 1 TJ, n=93 capillaries with 2 TJ. Data shown as mean ± SEM. Data shown as mean ± SEM. **(C)** Endothelial thickness plotted as a function of lumen area. **p=0.0011 (two-sided). Pearson r = 0.2374. R squared = 0.05637. **(D)** Endothelial thickness plotted as a function of lumen diameter. ***p=0.0009 (two-sided). Pearson r =0.2430. R squared = 0.05903.

We further asked whether heterogeneity in capillary diameter was related to the extent of pericyte coverage. Pericyte coverage was measured as the percentage of the capillary wall that was contacted by pericyte processes (**Supplementary Fig. 3**). We found no difference in pericyte coverage between 1 and 2 TJ capillaries.

Finally, we asked whether tissue fixation and processing could affect capillary diameter ranges in the Cortical MM^3 data. Insufficient intravascular pressure during transcranial perfusion and fixation procedures can lead to collapsed or altered vascular lumen within volume EM data. To collect ground truth data, in vivo deep two-photon imaging was performed in the visual cortex of 3 separate adult mice to measure capillary diameters from the pial surface to the gray and white matter interface (**Fig. 5A-C**). The capillary zone was defined identically as with volume EM quantification and measurements were distributed across all cortical layers. The diameter of capillaries in Cortical MM^3 were, on average, slightly smaller in diameter than that seen in vivo (**Fig. 5D**). However, the range of capillary diameters (∼3 to 5 μm) was similar between in vivo and volume EM data (**Fig. 5E**). This confirms that perfusion fixation and tissue handling used to generate the Cortical MM^3 data preserved the expected range in capillary diameter typically seen in vivo.

**FIGURE 5.**
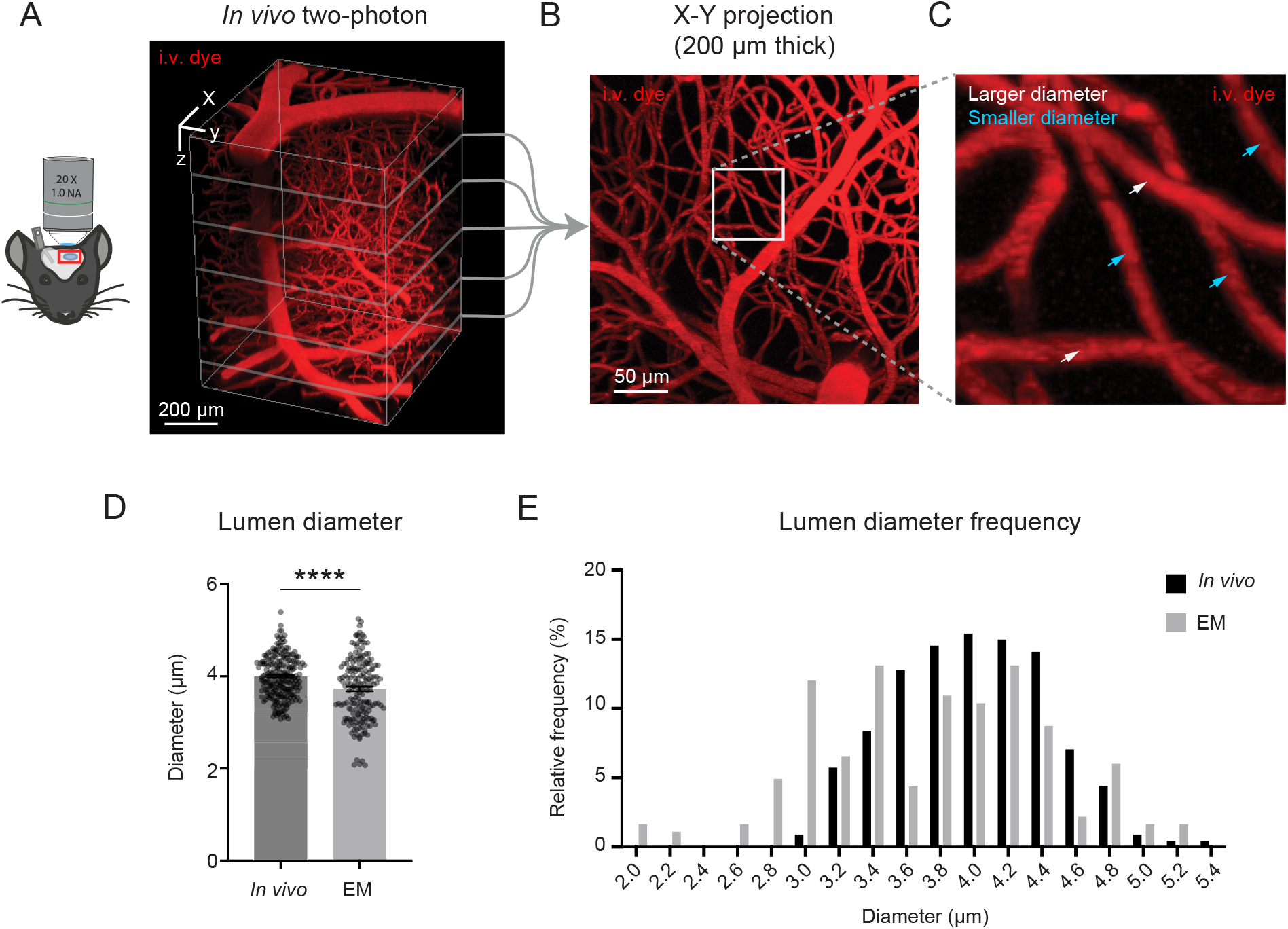
Volume EM data exhibits slightly smaller average capillary diameter than seen in vivo but retains heterogeneity in capillary diameter. **(A)** Deep in vivo two-photon imaging of isoflurane anesthetized mice via cranial window using Alexa 680-dextran (i.v. dye). 3D rendering of microvasculature within the mouse primary visual cortex. **(B**,**C)** Maximal projection from 250-450 μm of cortical depth. Inset shows example region of diameter measurement for individual capillaries, with white arrows showing larger capillaries and cyan arrows showing smaller capillaries. Capillary diameters were measured across all cortical layers. **(D)** Comparison of capillary lumen diameters between in vivo two-photon imaging and volume EM data. Unpaired t-test with Welch’s correction (two-sided), t(307.9)=4.670;****p<0.0001. N=227 capillaries from 3 adult mice for in vivo data; n=183 capillaries from 1 mouse for the MICrONS Cortical MM^3 data. Data shown as mean ± SEM. **(E)** Frequency distribution of lumen diameters from each data type.

## DISCUSSION

In this study, we used large-scale volume EM resources (20) to show that endothelial cell number has an influence on microvascular lumen size. Microvessels in transitional zones near penetrating arterioles and ascending venules are larger and typically composed of 2 to 5 endothelial cells, while the walls of capillaries intervening these regions are constructed from either 1 or 2 endothelial cells at a nearly 50:50 ratio. We show that capillaries with 2 endothelial cells are, on average, modestly larger than those with one endothelial cell. However, the diameter ranges of capillaries with 1 and 2 endothelial cells remains broad and heavily overlapping, suggesting additional mechanisms involved in establishing capillary diameter heterogeneity.

Endothelial architecture, such as cell density, is established during cerebrovascular development (22), and shaped by blood flow (23) and the metabolic demands (24) of the growing brain. Whether endothelial structure is stable in the adult brain or actively modeled has not been deeply examined. Some studies have tracked endothelial cells (and pericytes) longitudinally using in vivo two-photon imaging and showed relatively stable endothelial cell position and tight junction arrangement during adulthood, at least over the timeframe of days to weeks (25, 26). Cudmore et al. used a Tie2-based Cre driver to track capillaries in the motor cortex over time while mice had access to a running wheel (25). Interestingly, they reported marked stability of pericytes and endothelial cell density, except for small occasional shifts in nuclei position along the vessel wall. Reeson et al. showed that endothelial cells can be induced to reposition in the adult brain during local regression events caused by microvascular occlusions, indicating the potential to remodel in response to pathological stimuli (27). Endothelial cells that regress, migrate to nearby vessel segments and therefore increase endothelial cell number. Thus, contributions of endothelial structure to flow heterogeneity are likely established during development and stabilized during adulthood, yet remain responsive to flow perturbations in disease.

Our findings support the existence of additional mechanisms, beyond endothelial number, that control capillary diameter. As discussed above, capillary tone provided by capillary pericytes is a logical mechanism. There is now ample evidence for the contractile ability of capillary pericytes (28), but the endogenous vasoconstrictive signals that create this tone under basal conditions remain to be determined. Unraveling this mechanism will require experiments to conditionally delete receptors for vasoconstrictive signals known to be used by pericytes such as endothelin, thromboxane and noradrenaline receptors. Another potential contributor that remains poorly understood is tone generation from the cytoskeletal elements of the endothelial cells themselves, which is far less studied than pericyte contractility (29, 30). Further structural attributes of the vessel wall could include endothelial nuclei can protrude into the luminal space or the prevalence of small finger-like protrusions in the lumen called endothelial microvilli, which could both lead to increased local flow resistance (21).

There are some limitations to our study. First, the ultrastructural data is derived from the primary visual cortex from a single mouse. As volume EM data becomes more readily available, it will be possible to re-examine our hypothesis more broadly. Second, despite being able to sample hundreds of capillaries within Cortical MM^3, we only quantified ones cut perpendicular to the plane of highest spatial resolution, and therefore introduced bias toward a subset of capillaries oriented in one plane. Third, endothelial tight junction structure pericyte coverage varies along the longitudinally axis of the capillary segment. However, our analyses focused on 2D cross-sections of small regions along the longitudinal axis. Future analysis should perform 3D reconstructions of pericyte coverage, which will involve further proof-reading efforts to rigorously segment pericyte and endothelial compartments. Fourth, the issue of how tissue fixation and handling affect the native structure of the vascular lumen and wall components requires deeper studies. Fixation approaches have a strong influence when preserving the extracellular space during EM (31, 32). By comparing variance on capillary diameter between Cortical MM^3 and data collected from anesthetized mice using in vivo deep two-photon imaging, we see a similar range of capillary diameters, which lends confidence to the idea that capillary diameters were generally preserved.

In the Alzheimer’s brain, and in many related neurological diseases, capillary diameter is altered and microvascular density is reduced. This is expected to impair blood flow by increasing flow resistance but will also disrupt the range of capillary diameters that is critical for blood distribution. Further, shifts toward increased basal capillary heterogeneity may raise the threshold to distribute blood and oxygen during functional hyperemia. How alterations in endothelial structure and density factor into these disease-related microvascular deficits remains heavily understudied, yet vital to understanding mechanistic targets for improvement of microvascular perfusion.

## METHODS

### Volume EM data

#### Data collection from Cortical MM^3

Vasculature within the mouse visual cortex was identified in the publicly accessible volume EM resource (Cortical MM^3) created as part of the IARPA Machine Intelligence from Cortical Network (MICrONS) consortium project (https://www.microns-explorer.org/cortical-mm3). The data set included a 1.4 mm x .87 mm x .84 mm tissue volume from the visual cortex of a P87 mouse. Microvascular data was collected from all 6 cortical layers (and superficial callosal white matter) throughout the data set; n=179 vessels from gray matter and n=6 vessels from white matter. Care was taken not to sample from the same vessel segment twice. Vessels were identified as capillaries when they were branch order ≥ 4 from penetrating arterioles and ascending venules (both denoted as 0 order). Transition zone vessels ranged from branch orders 1-4 from penetrating arterioles (arteriole-capillary transition; ACT) or ascending venules (capillary-venous transition; CVT). To confirm vessels were branching from the arteriole side, ensheathing pericytes and perivascular fibroblasts were both required for ACT identity (33). To reduce variation in vessel and endothelial area, images/measurements were taken from areas without endothelial or pericyte nuclei present. This was more challenging within ACT and CVT zones, as nuclei were more abundant in this region. However, measurements from ACT zones were sometimes taken when the nuclei of perivascular fibroblasts was present.

#### EM data analysis

Screen captures of identified vessels were taken in Neuroglancer from the 2D EM view of the dataset with scale bars. Images of vessels were imported into and measured using FIJI/ImageJ (NIH), with the scale set according to individual scale bar provided in Neuroglancer. Lumen area/circumference and vessel area/circumference were measured using the freehand selection tool. Lumen diameter of capillary vessels, which were typically circular, was calculated as: SQRT(lumen area/pi). Due to the occasionally elliptical shapes of transition zone vessels, particularly on the venule side, the lumen diameter was measured using the straight-line tool on the minor axis. The endothelium area was calculated from the difference in vessel and lumen area. Endothelium thickness was measured with the straight-line tool at 5 different points around the vessel and averaged. Percent pericyte coverage was calculated by the length of the vessel circumference in contact with pericyte processes.

### In vivo imaging

#### Mice

In vivo deep two-photon imaging data from 3 adult mice was used for microvascular diameter measurements. The genotypes of these mice were Thy1-YFP (Jax: 003782; 1 mouse, 24 months old, male) and Atp13a5-2A-CreERT2-IRES-tdTomato (2 mice, 4 months old, male)(34), and all were on a C57Bl/6 background. Room temperature and humidity were maintained within 68–79□°F (setpoint 73□°F) and 30–70% (setpoint 50%), respectively. Mouse chow (LabDiet PicoLab 5053 irradiated diet for standard mice, and LabDiet PicoLab 5058 irradiated diet for breeders) was provided ad libitum. The Institutional Animal Care and Use Committee at the Seattle Children’s Research Institute approved all procedures used in this study (protocol #IACUC00396).

#### Surgery

Chronic cranial windows (skull removed, dura intact) were implanted in the skulls of all mice. Briefly, surgical plane anesthesia was induced with a cocktail consisting of fentanyl citrate (0.05□mg/kg), midazolam (5□mg/kg) and dexmedetomidine hydrochloride (0.5□mg/kg)(all from Patterson Veterinary). Dexamethasone (40 µL; Patterson Veterinary) was given 4–6□h prior to surgery to reduce brain swelling during the craniotomy. Circular craniotomies ∼4□mm in diameter were generated under sterile conditions and sealed with a glass coverslip consisting of a round 4□mm glass coverslip (Warner Instruments; 64-0724 (CS-4R)) glued to a round 5□mm coverslip (Warner Instruments; 64-0700 (CS-5R)) with UV-cured optical glue (Norland Products; 7110). The coverslip was positioned with the 4□mm side placed directly over the craniotomy, while the 5□mm coverslip laid on the skull surface at the edges of the craniotomy. An instant adhesive (Loctite Instant Adhesive 495) was carefully dispensed along the edge of the 5□mm coverslip to secure it to the skull. Lastly, the area around the cranial window was sealed with dental cement. This two-coverslip “plug” fits precisely into the craniotomy and helps to inhibit skull regrowth, thereby preserving the optical clarity of the window over months. Mice recovered for a minimum of 3 weeks following surgery.

#### Two-photon imaging

In vivo two-photon imaging system was performed with a Bruker Investigator (run by Prairie View 5.5 software) coupled to a Spectra-Physics Insight X3 laser source. Far red fluorescence emission was collected through a 700/75□nm bandpass filter, respectively, and detected by gallium arsenide phosphide photomultiplier tubes. Low-resolution maps of the cranial window were first collected for navigational purposes using a 4-X (0.16 NA) objective (Olympus; UPlanSAPO). We then switched to a 20-X (1.0 NA) water-immersion objective (Olympus; XLUMPLFLN) and used 1210□nm excitation to visualize the vasculature using intravenously injected Alexa-680 dextran, which was custom-conjugated following prior studies (35). All imaging with the water-immersion lens was done with room temperature distilled water. Imaging was performed generally over the primary visual cortex.

#### Quantification of vascular diameter from in vivo data

Capillary and transition zone vasculature were measured with the FIJI/ImageJ macro VasoMetrics (36), which provides the average diameter along the vessel segment based on full-width at half-max fluorescent intensity collected across multiple evenly distributed regions along the vessel length. As with the volume EM data, vascular metrics were acquired from all 6 cortical layers (and some callosal white matter) throughout the data set

#### Statistical Analysis

All statistical analysis was performed using GraphPad Prism v9. For all unpaired t-tests performed, normality and F tests for variance were performed. When statistically significant F tests were observed, unpaired t-test with Welch’s correction for unequal variances was performed. When data with non-normal Gaussian distributions were observed, nonparametric Mann-Whitney U tests were performed. Kruskal-Wallis tests were performed for analysis across CVT, capillary, and CVT zones.

## Declaration of interests

The authors have no conflict of interest that could be perceived as prejudicing the impartiality of the research reported.

## Funding

Our work is supported by grants to A. Shih from the NIH/NINDS (NS097775) and NIH/NIA (AG062738, R21AG069375, RF1AG077731).

## Author contribution statement

This project was conceived by AYS, and all analyses were performed by SMS. SKB and VCS served as independent raters of tight junction number analysis. YL and SS contributed in vivo deep imaging data sets. MT provided consultation on the MICrONS data set. The manuscript was written by AYS with feedback from all authors.

## Notes

### Competing Interest Statement

The authors have declared no competing interest.

