## Supplementary Figures for "Endothelial structure contributes to heterogeneity in brain capillary diameter"

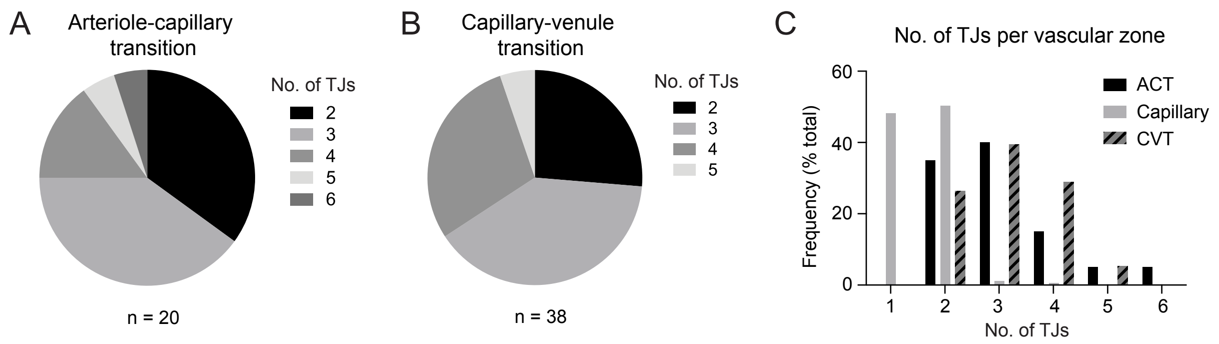


**Supplementary Figure 1.** Tight junction number in transitional zones. **(A,B)** Distribution of TJ number in ACT (A) and CVT (B) zones. **(C)** Frequency distribution of TJ number across ACT, capillary and CVT zones.

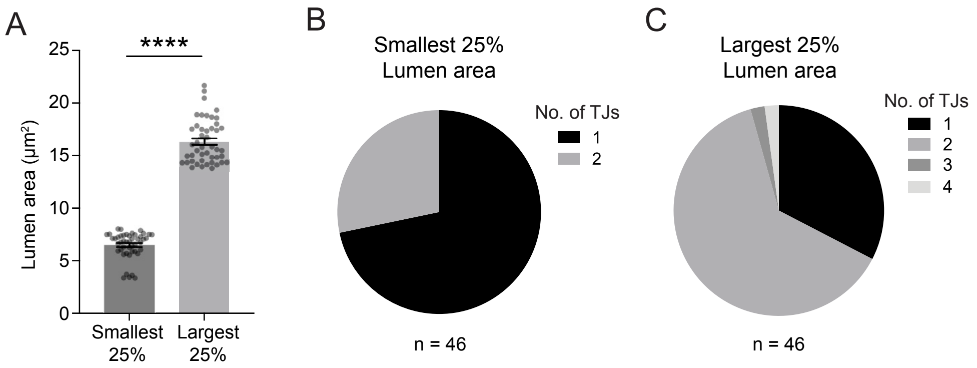


**Supplementary Figure 2.** Tight junction number in smallest and largest capillaries. **(A)** Lumen area for smallest and largest capillaries. Unpaired t test with Welch's correction (two-sided), t(73.66)=27.49;****p<0.0001. **(B,C)** Distribution of TJ number in the smallest and largest capillaries.

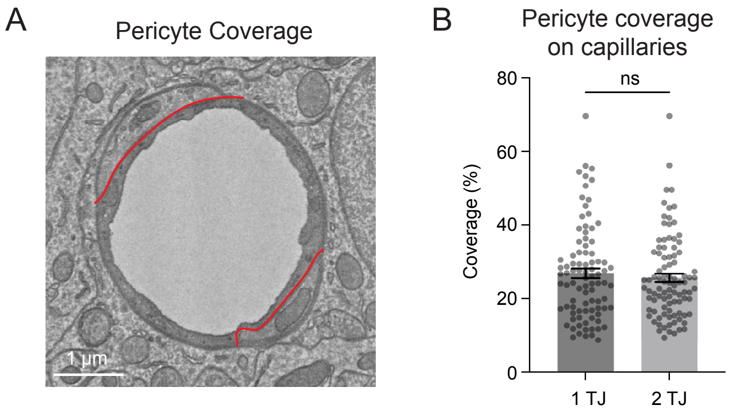


**Supplementary Figure 3.** Pericyte coverage between 1 and 2 TJ capillaries. **(A)** Example of capillary cross section with red lines marking the pericyte-endothelial interface. **(B)** Pericyte coverage between 1 TJ and 2 TJ capillaries. Unpaired t test (two-sided), t(180)=0.6902;p=0.4909. N=89 capillaries with 1 TJ, n=93 capillaries with 2 TJ. Data shown as mean ± SEM.
